# Heard or understood? Neural tracking of language features in a comprehensible story, an incomprehensible story and a word list

**DOI:** 10.1101/2022.11.22.517477

**Authors:** Marlies Gillis, Jonas Vanthornhout, Tom Francart

## Abstract

Speech comprehension is a complex neural process on which relies on activation and integration of multiple brain regions. In the current study, we evaluated whether speech comprehension can be investigated by neural tracking. Neural tracking is the phenomenon in which the brain responses time-lock to the rhythm of specific features in continuous speech. These features can be acoustic, i.e., acoustic tracking, or derived from the content of the speech using language properties, i.e., language tracking. We evaluated whether neural tracking of speech differs between a comprehensible story, an incomprehensible story, and a word list. We evaluated the neural responses to speech of 19 participants (6 men). No significant difference regarding acoustic tracking was found. However, significant language tracking was only found for the comprehensible story. The most prominent effect was visible to word surprisal, a language feature at the word level. The neural response to word surprisal showed a prominent negativity between 300 and 400 ms, similar to the N400 in evoked response paradigms. This N400 was significantly more negative when the story was comprehended, i.e., when words could be integrated in the context of previous words. These results show that language tracking can capture the effect of speech comprehension.

**Significance statement:** Most neural tracking studies focus on how the brain tracks acoustic speech features. However, whether acoustic tracking reflects speech comprehension is unclear. Therefore, in the pursuit of finding a neural marker for speech comprehension, language tracking might be a more suitable candidate. The results of this study showed that, indeed, language tracking can capture the effect of speech comprehension. This indicates that language tracking is a potential neural marker of speech comprehension. Such a neural marker would allow testing speech comprehension in populations that are currently difficult to test with behavioral tests, such as young children and persons with cognitive impairment.

## 1 Introduction

Speech comprehension is a complex neural process that relies on the activation and integration of different brain regions. Bottom-up and top-down processes allow speech comprehension to be fast and efficient, which is necessary to engage in conversations. However, the neural processes underlying speech comprehension are not well understood. Although fMRI can gain useful insight into these processes, with this neuro-imaging modality it is challenging to investigate the fast temporal responses to a rapid and dynamic speech signal. To overcome this, EEG or MEG can be used to investigate these responses with a higher temporal resolution, however compromising with a lower spatial resolution. To investigate the neural responses to sounds and speech, evoked responses to repetitive simple sounds can be investigated, for example, auditory brainstem response (for a review on evoked responses, see Picton, 2010). However, by using simple sounds, the insight gained is limited as these simple sounds do not require in-depth processing to yield comprehension. In recent years, there has been progress towards understanding the brain responses to continuous, natural running speech, relying on neural tracking. This is the phenomenon in which the brain time-locks to the rhythm of specific characteristics of the continuous speech (for a review, see Brodbeck and Simon, 2020). These characteristics can range from acoustic to language properties of speech.

Most neural tracking research focuses on acoustic aspects of the incoming sounds, called acoustic tracking. The link between acoustic tracking and speech comprehension is not clear. Some studies showed that, by using acoustic tracking of different speech fragments with varying signal-to-noise ratios, an objective measure could be derived similarly to the behaviorally measured speech comprehension (Vanthornhout et al., 2018; Lesenfants et al., 2019). This suggests that there is a link between acoustic tracking and speech comprehension. However, recent studies by Kosem et al. (2022) and Verschueren et al. (2022) showed that this link between acoustic tracking and comprehension is not always observed. More specifically, by manipulating speech comprehension by either using noise-vocoded speech or changing the speech rate, they observed that as the speech became comprehensible, acoustic tracking did not increase. Altogether, this suggests that acoustic tracking is required to comprehend speech, but it may not provide sufficient evidence to draw conclusions about the comprehension of speech. We might overcome this issue by investigating neural tracking of language properties of the speech, hereafter called language tracking (Gillis et al., 2022).

To measure language tracking, typically, an encoding modeling approach is used (Gillis et al., 2022). This approach yields two valuable outcomes: (1) a prediction accuracy which expresses *how well* the speech feature is represented in the brain responses, and (2) a temporal response function (TRF), which denotes *how* the brain responds to the speech feature. The encoding modeling approach can be applied using any speech characteristic which represents the speech over time. Previous studies often focussed on acoustic features, which represent the acoustic energy of the speech signal. In order to measure language tracking, language features are used which are derived from the content of the speech signal. Such language features characterize the speech signal as sparse arrays with an impulse at the onset of a phoneme or word. The amplitude of the impulse is weighted by a specific language property of the considered phoneme or word, e.g., surprisal.

Over the last few years, language tracking has attracted more attention. Initially, it was identified which language features were significantly tracked by the brain (e.g. Broderick et al., 2018; Brodbeck et al., 2018; Weissbart et al., 2020; Gillis et al., 2021; Heilbron et al., 2022). Language tracking was seen at the level of phonemes, showing negative responses around 250 ms to 300 ms (further denoted as *N* 250*_T RF_*) for EEG (Gillis et al., 2021; Di Liberto et al., 2019). At the level of words, studies showed that the brain shows a negative response around 400 ms after word onset (further denoted as *N* 400*_T RF_*), which is modulated by the information conveyed by the word, indicated by, e.g., word surprisal (Weissbart et al., 2020). This response is assumed to be similar to the N400 as observed in evoked response studies and it is thought to resemble the activation of the lexical item and integration of this item into the context of the previous words (for a review on the N400, see Lau et al., 2008; Kutas and Federmeier, 2011). More recently, it was assessed whether language tracking is related to behavioral speech comprehension. By comparing listeners listening to their second language to native speakers of the language, Di Liberto et al. (2021) showed that as language proficiency increases, the response around 400 ms becomes more similar to a native speaker. However, their study relied on a different group of participants for each language proficiency while group size and age distribution differed across groups. Keeping the group of participants consistent, two other studies manipulated the amount of speech comprehension by manipulating the speech via word scrambling (Broderick et al., 2020) or manipulating the speech rate (Verschueren et al., 2022). Both studies observed that as the speech became incomprehensible, the *N* 400*_T RF_* disappeared. However, these studies did not rely on natural spoken speech as they relied on either synthetic scrambled speech (Broderick et al., 2020) or artificially sped up speech (Verschueren et al., 2022).

In this study, we used natural spoken speech. We presented three different speech materials, a comprehensible story, an incomprehensible story and a word list, naturally spoken by the same speaker and keeping the group of participants consistent. We hypothesized that when native Flemish participants listened to a story in a foreign language, here Frisian, they did not track the content of the presented speech, and thus we would not see language tracking, implying that language features would not improve the prediction accuracy nor a *N* 250*_T RF_* or *N* 400*_T RF_* would be observed. For the word list, the listener does have access to the meaning of the word, but the word cannot be integrated in the context of the previous words. Therefore, both processes would affect the presence of the *N* 400*_T RF_* to language features at the word level. While language features at the phoneme level, which elicit an *N* 250*_T RF_*, could be robust as phoneme probabilities are not changed. Overall, this study aims to assess whether measures of language tracking can capture the effect of speech comprehension.

## 2 Methods

### 2.1 Participants

For this study, 19 participants were included (6 men; mean age *±* std = 22 *±* 3 years). The inclusion criteria were normal hearing, Dutch as a mother tongue and the absence of attention problems, learning disabilities, or severe head trauma. The latter were identified via a questionnaire. Pure tone audiometry was conducted at center frequencies from 125 to 8000 Hz to assess the hearing capacity. Participants, where a hearing threshold exceeded 20 dB HL, were excluded from this study.

### 2.2 Speech stimuli

We created custom speech stimuli for this study, resulting in three conditions: Dutch speech, Frisian speech and a word list, all spoken by the same speaker. The speech materials were derived from two episodes of ‘De Volksjury’, a Flemish podcast series about crime cases. The podcast episodes were transcribed and rewritten in an entertaining story, resulting in two fragments: a story about Jack The Ripper and a shorter story about the disappearance of Madeleine McCann. The latter consisted of only the beginning part of the original podcast and was used to create the Frisian story and the Dutch word list. To create the Frisian story, the story was translated into Frisian. We opted for Frisian as this language is related to Dutch and follows similar grammatical rules. However, it is poorly understood by Dutch listeners without knowledge of Frisian. For the Dutch word list, all the words in the story were randomly shuffled, thereby not preserving the syntax. The word list contained sentences matched in length with the original story to aid the speaker in using natural prosody.

The speech materials were all spoken by the same native Dutch male speaker who learned Frisian as a second language. We manually edited the speech recordings to remove stutters, clicks, and mispronunciations. In the Frisian story, we removed sentences that consisted of a high amount of loanwords. In total 5 sentences, i.e. 52 words or around 4% of the words, were removed from the speech material. After inspection, for the word list, we removed sentences with unnatural prosody, including odd pauses and intonation (in total 199 words, i.e., 14% of the words).

The Dutch story lasted around 45 minutes and was divided in four parts, each lasting between 10 to 13 minutes. Only the first part of this speech material is presented in the current study, matched to the length of the other speech materials.

The Frisian story lasted around 7 minutes, and the word list around 9 minutes. As the listener could not understand these two speech materials, we aimed to motivate the listener to attend attentively to the stimulus by keeping the speech material shorter and making the listener perform a word identification task. Therefore, the latter two speech materials were divided in four parts, each lasting around 2 minutes. Following every 2-minute segment, the listener was prompted with three words and required to indicate whether each word was presented in the preceding 2-minute fragment.

The speech was presented bilaterally to the participants at an intensity of 65 dB (A-weighted). The speech materials were presented in a different random order for each participant; however, the order of the fragments within one speech material was preserved. After each fragment, a content question was asked in case of the Dutch speech fragments, or a word identification task was conducted to keep the participant motivated to listen to the stimulus attentively. Additionally, we asked the participant to rate their speech understanding by asking the question *how much did you understand about the content of the story?* ranging from 0 to 100%. Note that even though individual words could be understood, the content of the story is not necessarily understood, as was the case for the Dutch word list.

### 2.3 Speech features

Similar to Gillis et al. (2021), we used 8 speech features. To control for the acoustic energy of the speech signal, we used two continuous speech features, i.e., the speech spectrogram and acoustic edges. We included two (pre-)lexical speech features which denote the onset of phonemes and words. Although neural responses to these (pre-)lexical speech features are not purely acoustic, these features are included in the model as acoustic controls for the language features. Additionally, we included four language features, i.e., phoneme surprisal, cohort entropy, word surprisal and word frequency. Unlike the acoustic and (pre-)lexical features, the language features are derived from the content of the speech. All speech features for the first sentence of the three speech materials are visualized in Figure 1.

**Figure 1:**
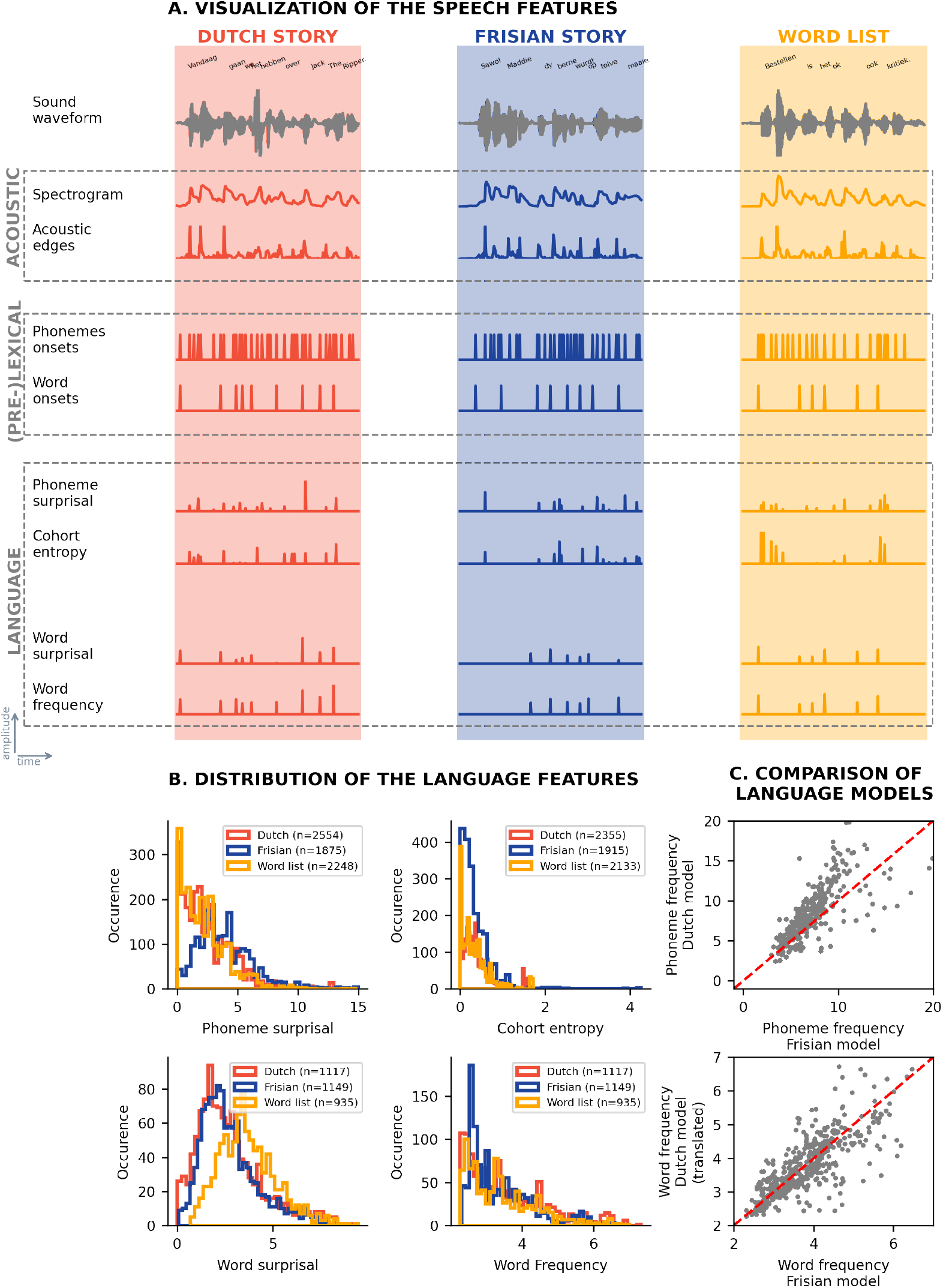
Speech features at acoustic, (pre-)lexical and language level. We visualized the speech features used in this study for the three speech materials. In panel A, the different features are visualized for the first sentence of each speech material, respectively the Dutch story (orange), the Frisian story (blue) and the word list (yellow). In panel B, the distribution of the non-zero values of the different language features is depicted. In panel C, the Dutch and Frisian language models are compared. For the top plot, the phoneme frequency at a specific place in the word in Dutch compared to Frisian is visualized. In the plot below, the word frequencies are visualized for the Frisian word, derived from the Frisian language model, versus its Dutch translation, derived from the Dutch language model.

The spectrogram and acoustic edges reflect the continuous acoustic energy of the speech material. The spectrogram was created using the Gammatone Filterbank Toolkit 1.0 (Heeris, 2014, center frequencies between 70 and 4000 Hz with 256 filter channels and an integration window of 0.01 second). By combining 32 consecutive filter outputs, the 256 filter outputs were averaged in eight frequency bands. Additionally, the spectrogram was downsampled to match the sampling frequency of the preprocessed electroencephalography (EEG) data, namely 128 Hz. To obtain the acoustic edges, a half-wave rectification was applied to the derivative of the spectrogram, i.e., negative values of the derivative were clipped 0.

Using a forced aligner (Duchateau et al., 2009), we constructed a word and phoneme segmentation. This forced aligner was also applied for the Frisian condition and corrected manually with the PRAAT software (Boersma and Van Heuven, 2001). These segmentations were used to create the (pre-)lexical and language features. All these features are discrete, one-dimensional arrays with impulses on onsets of phonemes or words. These impulses have a fixed amplitude for the (pre-)lexical features, while for the language features, the impulses are modulated by the information value of a phoneme or word.

Phoneme surprisal and cohort entropy are two language features at the phoneme level. Both metrics are derived from the active cohort of words, i.e., the words congruent with the previously uttered phonemes. Phoneme surprisal reflects how surprising a phoneme is, given its previously uttered phonemes. It reflects the phoneme’s prediction error, which is intrinsically related to the information a phoneme contributes to its context. The more likely a specific phoneme is, the lower its surprisal value, and the less information is gained (and vice versa). Cohort entropy reflects the uncertainty of the phoneme prediction. If the uncertainty of the phoneme prediction is high, the active cohort consists of a large number of words with a similar probability of being uttered.

Phoneme surprisal is calculated as a negative logarithm of the conditional probability of the phoneme given the previously uttered phonemes. Cohort entropy is calculated as the Shannon entropy of the active cohort of words. To project cohort entropy to similar scales, the values of cohort entropy were divided by the maximal value of cohort entropy in the speech fragment. More details and the mathematical determinations of these metrics can be found in Brodbeck et al. (2018). For the Dutch language features at the phoneme level, the phoneme segmentation of the words was obtained using a custom word-to-phoneme dictionary (9082 entries), while the word frequencies were derived from SUBTLEX-NL (Keuleers et al., 2010) (134724 entries). For the Frisian language features at the phoneme level, both the word-to-phoneme dictionary (75036 entries) and the word frequencies (124166 entries) were used from Yilmaz et al. (2016).

By applying the Kolmogorov-Smirnov test, we observed that the distribution of the phoneme surprisal

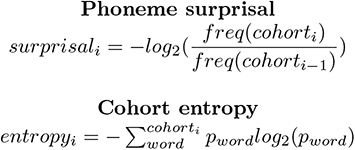

values in the Frisian condition is significantly different from the distribution of the values for the Dutch story and the word list. For cohort entropy, the distribution of significantly different between all three speech conditions. Additional details are included in the supplementary material (section 6.1).

Similar to phoneme surprisal, word surprisal reflects how surprising a word is given the previously uttered words. It reflects the word’s prediction error, which is related to how much information the word contributes to its context. Word frequency reflects the probability of the word independent of its context.

Word surprisal was calculated as the negative logarithm of the conditional probability of the considered word given the four preceding words. This conditional probability was obtained with a 5-gram model created by Verwimp et al. (2019) for the Dutch material and by Yilmaz et al. (2016) for the Frisian material. Using these models, word frequency was obtained by calculating the negative probability of the uni-gram probability of the word.

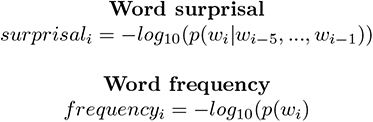

No significant differences were observed between the distributions of the three speech conditions for both word surprisal and word frequency. Additional details are included in the supplementary material (section 6.1).

The language models used to derive the Dutch and Frisian language features differ. Therefore, we investigated the phoneme and word frequency to assess whether the language model had a significant impact on the values of the speech features. In the first comparison, we compared the probability of a phoneme occurring at a specific position in Dutch to this probability in Frisian, e.g., the probability of the phoneme /a/ occurring second is fairly similar in Dutch as in Frisian (visualized in figure 1.C top panel). These phoneme frequencies in Dutch were significantly correlated with those in Frisian (Pearson’s r = 0.68, p *<* 0.001). This suggests that phoneme frequencies are similar in Dutch and Frisian. Additionally, we compared the word frequency of a Dutch word derived from the Dutch language model to the word frequency of its Frisian translation using the Frisian language model. Indeed, these word frequencies are significantly correlated (Pearson’s r = 0.88, p *<* 0.001). Using language models of different sizes, which are also trained on different text corpora, is sub-optimal. However, these outcomes reassure us that the usage of these different language models is valid to do.

### 2.4 EEG preprocessing

To decrease preprocessing time, the measured EEG data were downsampled to 265 Hz (using the *resample* function of MATLAB which includes anti-aliasing). After this initial downsampling step, the spatio-temporal characteristics of eye-blink artifacts were estimated in the EEG to the second part of the Dutch story, using the multichannel Wiener filtering (Somers et al., 2018). These characteristics are then used to filter out eye-blink artifacts in the EEG measured to the first part of the Dutch story, the Frisian story, and the word list. Subsequently, the EEG data were referenced to the common average reference. The EEG data and the continuous acoustic speech features were filtered between 0.5 and 25 Hz using Least Squares Filtering (using least-squares linear-phase FIR filter function of MATLAB with the high-pass filter order of 5000, low-pass filter order of 250, with a stopband frequency 10% outside the passband frequency and passband weight of 100 and stopband weight of 1). Afterwards, the EEG data were downsampled to 128 Hz (using the *resample* MATLAB function) to reduce the time to compute the encoding models. Additionally, the EEG data were normalized, i.e., z-scoring (subtracted the mean and divided by the standard deviation). This normalization was performed across the EEG data of the three conditions for each EEG channel separately.

As the amount of data is substantially lower compared to previous studies investigating language tracking, we opted to further denoise the data to improve the SNR of the stimulus-related responses. To do so, we opted to apply denoising source separation (DSS) (de Cheveigné and Simon, 2008). This method allows for further removal of non-stimulus-related activity. It estimates spatial filters and associated components which capture the stimulus-related activity repeated across the different trials or participants, sorted according to power. Here, we estimated the spatial filters across the EEG data of all participants listening to the second part of the Dutch story. To train DSS, we used a filter between 2 and 25 Hz (Least square filtering with order 400; see further filter descriptions above). Note that this filter band is only used to train the DSS filters. The high-pass frequency was increased to 2 Hz to decrease the power of the lower frequencies. Nevertheless, note that 2 Hz is an arbitrary choice. Additionally, this second part is solely used to estimate the DSS components and to estimate the filter for the multichannel Wiener filtering to prevent a biased analysis. Subsequently, we selected the six components with the highest power to apply to the first part of the Dutch story, the Frisian story and the word list. Using these six components, we projected the EEG data back to sensor space.

Using a limited amount of data, one can expect a higher performance of the model when more training data is used (Mesik and Wojtczak, 2022). To avoid this bias, we limited the duration to the first 455 seconds for all three speech materials (discarding most of the data in the Dutch condition).

### 2.5 Encoding modeling approach

We applied a linear encoding model to get insight in how the brain responds to different speech features. This modeling approach estimates the spatial and temporal pattern of the neural responses to the speech features. This pattern is called a TRF. This TRF can then be used to predict the EEG responses associated with specific speech features. A prediction accuracy can be obtained by correlating the predicted EEG responses with the measured EEG responses. The prediction accuracy is a measure of neural tracking: looking at the brain responses of one subject, the more the brain *tracks* the speech, the higher the prediction accuracy will be.

To apply this encoding modeling approach, we used the Eelbrain Toolbox (Brodbeck, 2020). This toolbox estimates the TRF using an iterative boosting approach (David et al., 2007). We estimate the TRF using 10-fold cross-validation. This implies that the data is divided in ten equally long folds: 8 folds are used for estimating the TRF, i.e., training the model, one fold for validation of TRF estimation, and one fold to evaluate this TRF by determining the prediction accuracy, i.e., testing the model. This is repeated for all train and test combinations. Further used parameters for the model estimation were an integration window from -100 to 600 ms, a convolution with a basis window of 50 ms (Hamming window) which was used to make the TRF less sparse and the use of early stopping based on the *ℓ*2-norm of the difference between the actual and predicted EEG responses.

For the spectrogram and acoustic edges, the TRF is visualized as an average TRF across the different frequency bands. We included the TRF for each frequency band as supplementary figures (Figures S.1 and S.2).

### 2.6 Determination of acoustic and language tracking

Acoustic tracking is here denoted as the prediction accuracy obtained with a model which includes acoustic speech features, i.e., the spectrogram and acoustic edges.

To determine whether or not the brain significantly tracked the language features, we used an approach that relies on the difference in prediction accuracies between two models (as applied in, e.g., Di Liberto et al., 2015; Brodbeck et al., 2018; Gillis et al., 2021, ; for a visual representation see Figure 2). Here, we focused on the added value of all four language features combined. We constructed a baseline model, which included the acoustic and (pre-)lexical features, which served as acoustic controls for the included language features. Subsequently, we constructed a complete model that included the language features on top of the acoustic and (pre-)lexical features. The increase in prediction accuracy between the complete and baseline model signifies the improvement of the model due to the language features. Similarly, to investigate whether (pre-)lexical features improve the acoustic model, the performance of the acoustic model was subtracted from the model including acoustic and (pre-)lexical features.

**Figure 2:**
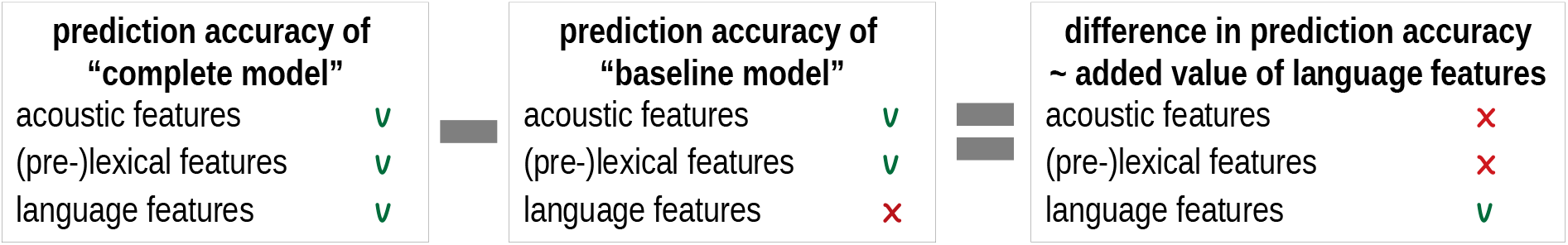
Methodological approach to determine the added value of language features. The added value of language features was determined as the difference between two models, respectively, the complete and baseline model. We considered (pre-)lexical features, denoting the onset of phonemes and words as control features and thus they are included in the baseline model. To determine the added value of language features, we considered all language features combined (i.e., phoneme surprisal, cohort entropy, word surprisal, and word frequency).

### 2.7 Determine the peak characteristics

We defined the maximal amplitude of the TRF, averaged across a channel selection, within specific time regions to determine the peak amplitudes and latencies. The time regions were determined based upon the average TRF of each speech feature. The channel selections were set a priori. For acoustic tracking, we used the channel selection suggested by Lesenfants et al. (2019) (F5, F3, F1, Fz, F2, F4, F6, FC5, FC1, FC3, FCz, FC2, FC4, FC6, C5, C3, C1, Cz, C2, C4, C6), while for language tracking, the same channel selection as Gillis et al. (2021) (P1, Pz, P2, CP1, CPz, CP2) was used. If the found peak was at the edges of the time region, the peak was discarded. The time regions used for the different peaks for each feature and the number of peaks are summarized in Table 1.

**Table 1:** Time regions selected per speech feature to determine the peak characteristics. The number of participants for whom a peak was found is reported as well (of the 19 participants included in this study).

| feature | time region | peak name | Dutch story | Frisian story | word list |
| --- | --- | --- | --- | --- | --- |
| spectrogram | 0 - 65 ms | P1 | 12 | 11 | 11 |
|  | 70 - 110 ms | N1 | 4 | 6 | 6 |
|  | 115 - 200 ms | P2 | 9 | 14 | 9 |
| acoustic edges | 0 - 90 ms | P1 | 19 | 18 | 18 |
|  | 95 - 140 ms | N1 | 12 | 7 | 9 |
| phoneme surprisal | 150 - 350 ms | $N250_{TRF}$ | 17 | 17 | 17 |
| cohort entropy | 150 - 350 ms | $N250_{TRF}$ | 14 | 18 | 18 |
| word surprisal | 300 - 500 ms | $N400_{TRF}$ | 17 | 16 | 17 |
| word frequency | 300 - 500 ms | $N400_{TRF}$ | 16 | 16 | 18 |

### 2.8 Statistics

We performed pairwise comparisons to compare whether the behavioral results, neural tracking, or neural response amplitudes and latencies differed between the different speech materials. As not all the data were normally distributed, we opted for non-parametric tests, namely the pairwise Wilcoxon signed-rank test and Mann-Whitney U Test, with a Benjamini-Hochberg correction for multiple comparisons. To investigate the differences between the speech materials for the behavioral results and for the neural tracking, we used the paired Wilcoxon signed-rank test, grouping the results from the same subject. However, these paired tests could not be used when investigating the differences in neural response amplitude or latency. For some subjects, no peak could be found, and the paired Wilcoxon sign-rank test cannot handle missing data. Therefore, the Mann-Whitney U Test was used in these cases. The outcomes of these tests are reported with the *z*- or *u*-score and p-value for, respectively, the Wilcoxon signed-rank test or Mann-Whitney U Test. These tests were applied using the Eelbrain (Brodbeck, 2020) implementation for pairwise comparisons.

To determine whether the TRF or the spatial distribution of the (increase in) prediction accuracy were significantly different from 0, we used cluster-based permutation tests as recommended by Maris and Oostenveld (2007), using the Eelbrain (Brodbeck, 2020) implementation. This test was applied for each TRF individually with a significance level of *α*=0.05 (by restriction of the threshold for forming clusters, i.e., *pmin*-argument in the Eelbrain implementation). Afterward, the resulting p-values per test were corrected for 3 comparisons using Benjamini-Hochberg correction for multiple comparisons.

## 3 Results

### 3.1 Behavioral Results

The behavioral results obtained in this study consisted of a subjective rating (*how much did you understand about the content of the story?*) and a word identification task which the participants only did for the Frisian story and the word list. The behavioral results are visualized in Figure 3.

**Figure 3:**
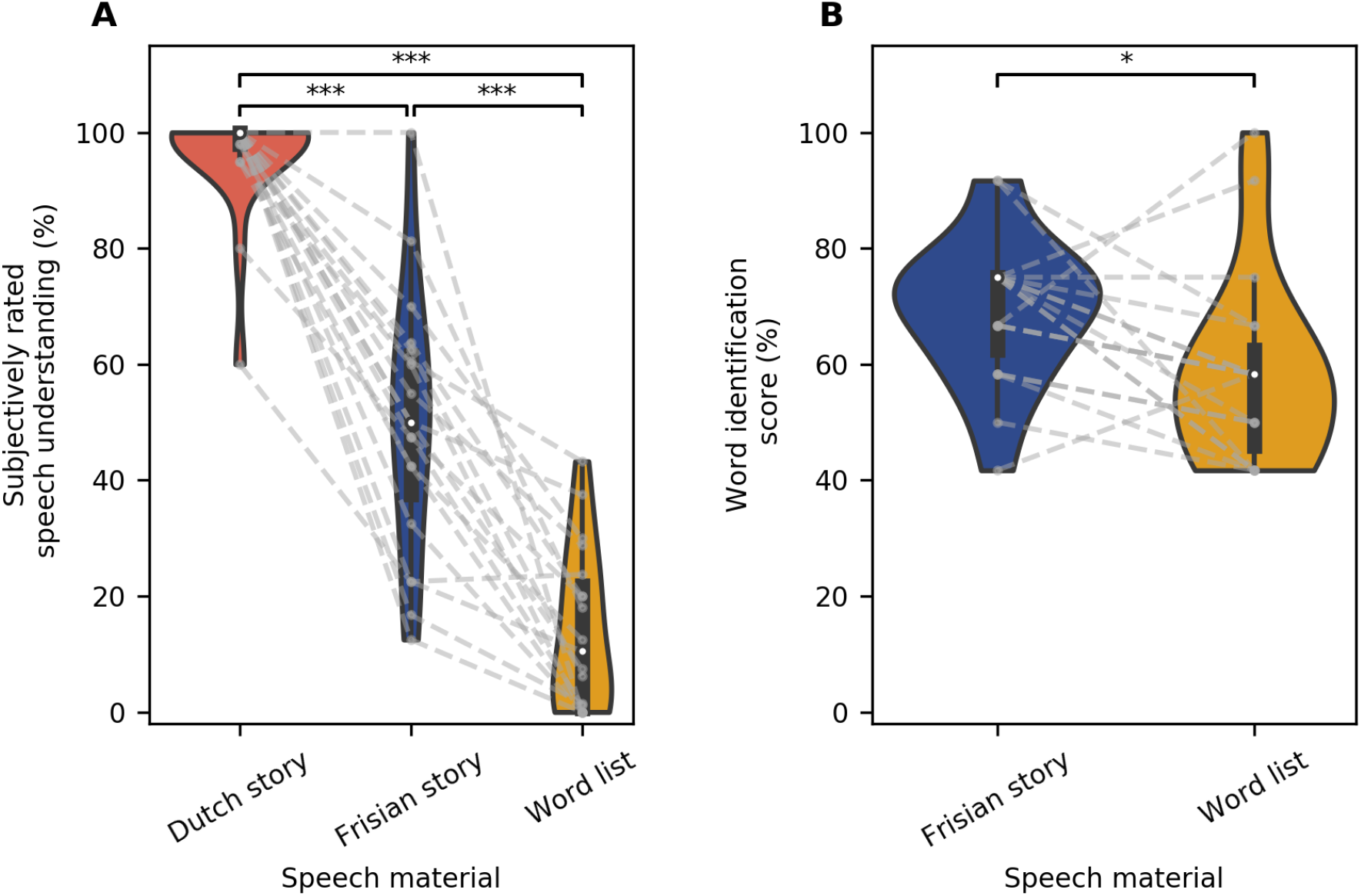
Behavioral results. Panel A: The subjectively rated speech understanding as a function of the speech material, respectively the Dutch story (orange), Frisian story (blue) and word list (yellow). Panel B: The results of the word identification task for the Frisian story and the word list. *(*: p < 0.05, **:p < 0.01, ***: p < 0.001)*

There are significant differences between the subjective rating of all three speech materials. The subjective ratings for the Dutch story (median = 100 %) were significantly different from those for Frisian (median = 50 %; z_(18)_ *<* 0.01, p *<* 0.01) and the word list (median = 10.5 %; z_(18)_ *<* 0.01, p *<* 0.01). Also, the subjective rating for the Frisian story differed significantly from the rating for the word list (z_(18)_ *<* 0.01, p *<* 0.01). These results show that the Dutch story is better comprehended than the Frisian story, which is better comprehended than the word list.

The results on the word identification score, were significantly higher for the Frisian story (median = 75 %) than the word list (median = 58.3 %; z_(18)_ = 35.50, p = 0.029).

### 3.2 Neural Tracking

To evaluate the acoustic tracking of speech, we investigated the prediction accuracies obtained with the acoustic model, i.e., the model including only the spectrogram and acoustic edges (shown in panel A of Figure 4). The results of the cluster-based permutation test showed that for all three speech materials, a prediction accuracy significantly higher than zero is obtained (1 cluster containing all 64 EEG channels, p *<* 0.001 for all speech materials; corrected for 3 comparisons). Additionally, we investigated the differences between the three speech materials by applying a pairwise comparison test on the prediction accuracies averaged across a frontal channel selection. The results of the pairwise comparison test did not identify a significant difference between the three speech materials.

**Figure 4:**
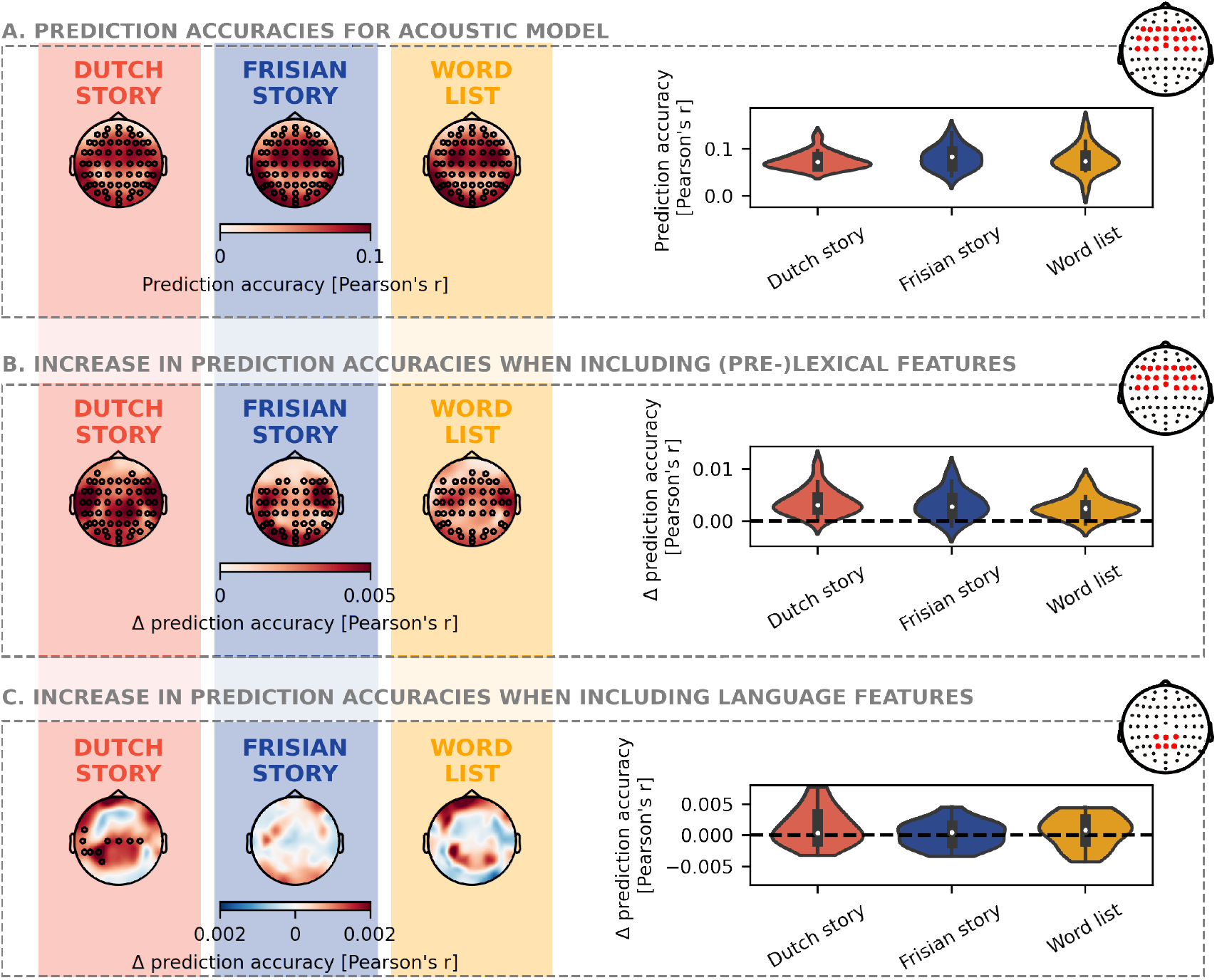
Neural tracking: Panel A: The prediction accuracies obtained with the acoustic model are shown for the three speech materials, respectively the Dutch story (orange), Frisian story (blue), and word list (yellow). More specifically, the spatial distribution of the prediction accuracies obtained with only the acoustic features is visualized on the left side. The same prediction accuracies are visualized on the right side but averaged across a frontal channel selection (see head plot). Panel B: The increase in prediction accuracies when including the (pre-)lexical features on top of the acoustic features are shown for the three speech materials. This increase in prediction accuracies, averaged across a frontal channel selection (see head plot), are visualized on the right side. Panel C: The increase in prediction accuracy when including the language features on top of a model with (pre-)lexical and acoustic features for all three speech materials. The same prediction accuracies are visualized on the right side but averaged across a central channel selection (see head plot). The black annotated channels indicate the cluster, which drives the spatial distribution to be significantly higher than zero as obtained by the cluster-based permutation test.

Additionally, we investigate whether the (pre-)lexical features significantly improve the prediction accuracy over acoustic features (shown in panel B of Figure 4). Indeed, for all three speech materials, one big cluster, covering occipital, temporal, and central areas, is observed which drove the increase in prediction accuracy to be significantly different from zero (all clusters obtained a cluster with p *<* 0.001; corrected for 3 comparisons). The pairwise comparisons of the increase in prediction accuracies averaged across a frontal channel selection did not show significant differences between the three speech materials.

Lastly, we investigated whether the prediction accuracies increase when language features are included on top of acoustic and (pre-)lexical features (shown in panel C of Figure 4). Only for the Dutch story, the results of the cluster-based permutation test showed that some central and left-lateralized channels drove the increase in prediction accuracy to be significantly different from zero (p = 0.045; corrected for 3 comparisons). However, the pairwise comparisons of the increase in prediction accuracies averaged across a central channel selection did not show significant differences between the three speech materials.

### 3.3 Neural response to acoustic and (pre-)lexical features

To investigate how the brain responds to acoustic speech features, one can investigate the pattern of the TRF (see Figure 5). The TRF to the spectrogram is characterized by three prominent peaks: a first positive peak (P1) followed a negative peak (N1) and a later second positive peak (P2). This pattern differs from the one seen for acoustic edges, where only a first positive peak (P1) and a following negative peak (N1) are observed. If a cluster was found within the time region of interest of the peak (with the same polarity as the expected polarity of the peak), we concluded that a peak was observed. Subsequently, we investigated whether there are differences in amplitude and latency of these peaks among the three different conditions.

**Figure 5:**
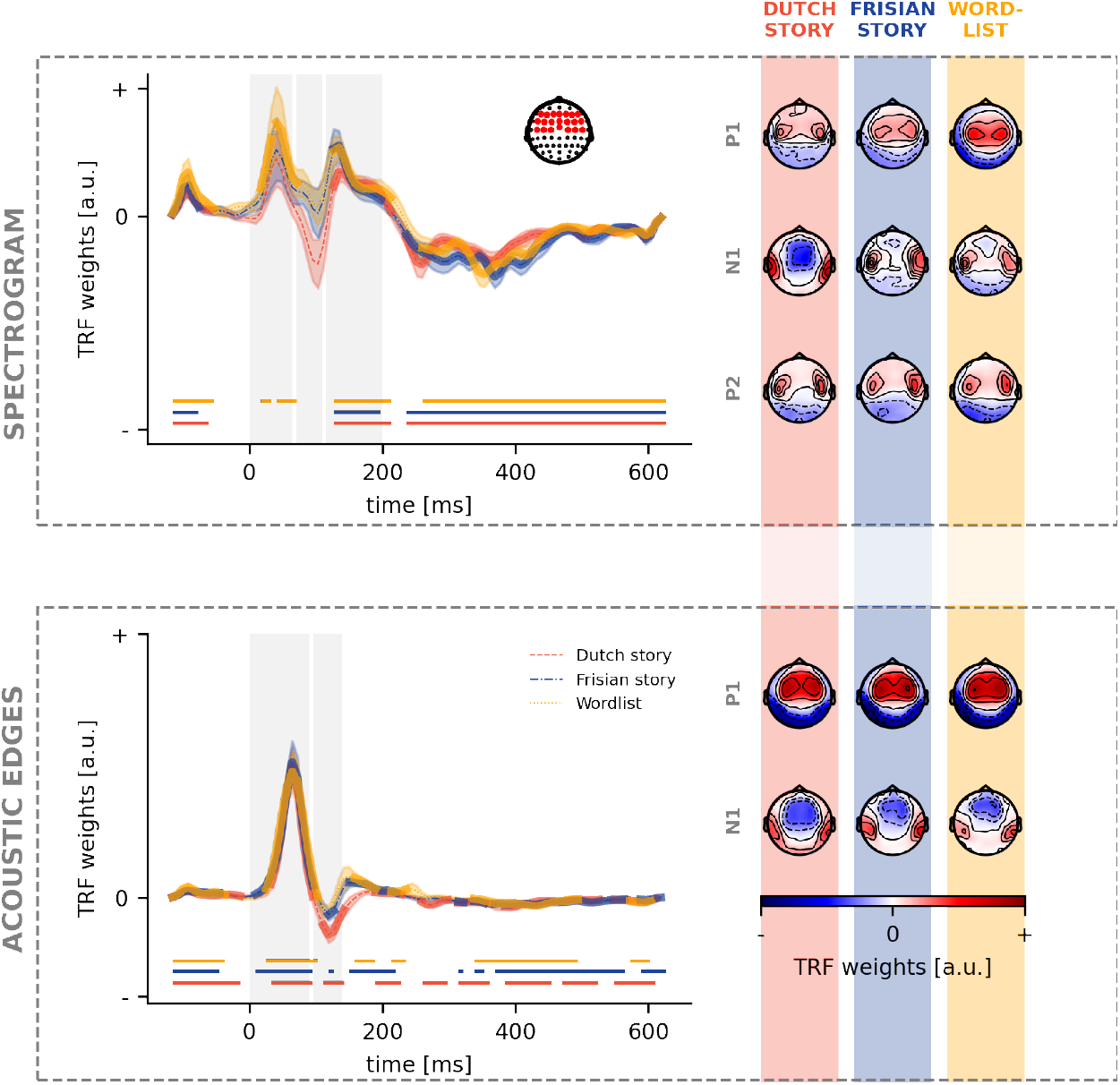
Neural response to acoustic speech features. For each acoustic speech feature, the TRFs in a frontal channel selection (shown as a right inset on the top panel) are depicted (left). The significance of the TRF is indicated by the thick lines in the corresponding colour and the horizontal lines below the TRF. The vertical grey blocks mark the time regions used for peak determination. For each of these peaks, the associated peak topographies are visualized (right) for respectively the Dutch story (orange), Frisian story (blue) and word list (yellow).

For the spectrogram, the P1 peak is observed for only the word list condition. No N1 peak for the spectrogram was observed for all three conditions, however, the P2 peak was for all conditions (top panel in Figure 5). At the same time, for acoustic edges, all conditions showed a P1 peak. However, only the Dutch story and the Frisian story showed a negative N1 peak (bottom panel in Figure 5). We investigated the peak amplitudes and latencies to quantify the differences between the TRFs. For all peak amplitudes and latencies, no significant differences were found between the three conditions for all peaks of the TRF to acoustic features.

Regarding phoneme onsets, we observe a P1 peak for all three conditions, while only an N1 peak for the Dutch story (top panel in Figure 6). No differences were observed regarding amplitude or latency among the three conditions. For word onsets, a P1 peak is observed for all three conditions, while none showed an N1 peak. The P2 peak was seen for all three conditions (bottom panel in Figure 6). For the P2 peak, we observed that the amplitude for the Frisian condition (median = 0.00037) was significantly different from the Dutch story (median = 0.00009; u_(28)_ = 222, p = 0.04; corrected after Benjamini-Hochberg), however, no differences were observed in the latency of the peaks.

**Figure 6:**
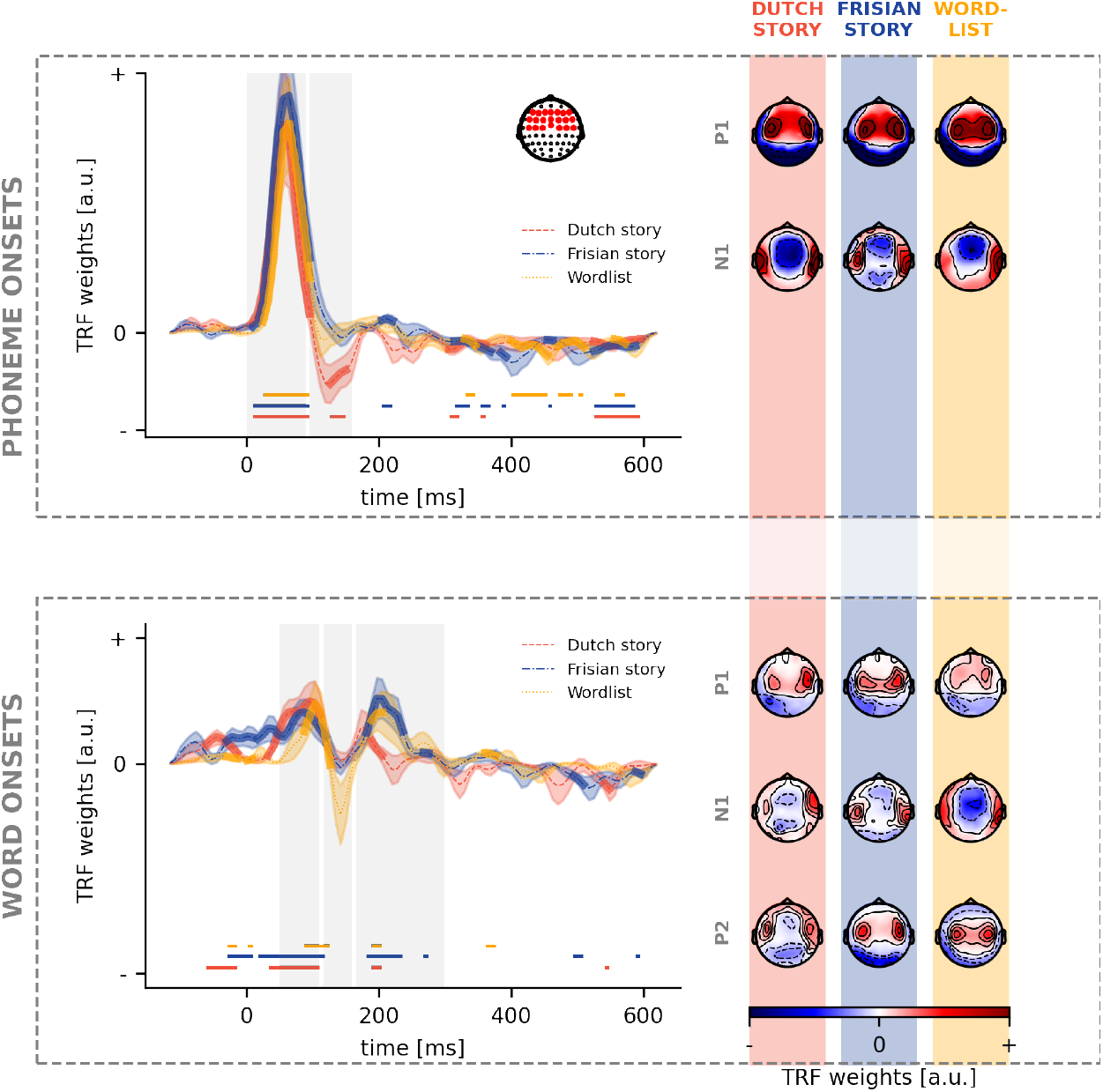
Neural response to (pre-)lexical speech features. For each (pre-)lexical speech feature, the TRFs in a frontal channel selection (shown as a right inset on the top panel) are depicted (left). The significance of the TRF is indicated by the thick lines in the corresponding colour and the horizontal lines below the TRF. The vertical grey blocks mark the time regions used for peak determination. For each of these peaks, the associated peak topographies are visualized (right) for respectively the Dutch story (orange), Frisian story (blue), and word list (yellow).

**Figure 7:**
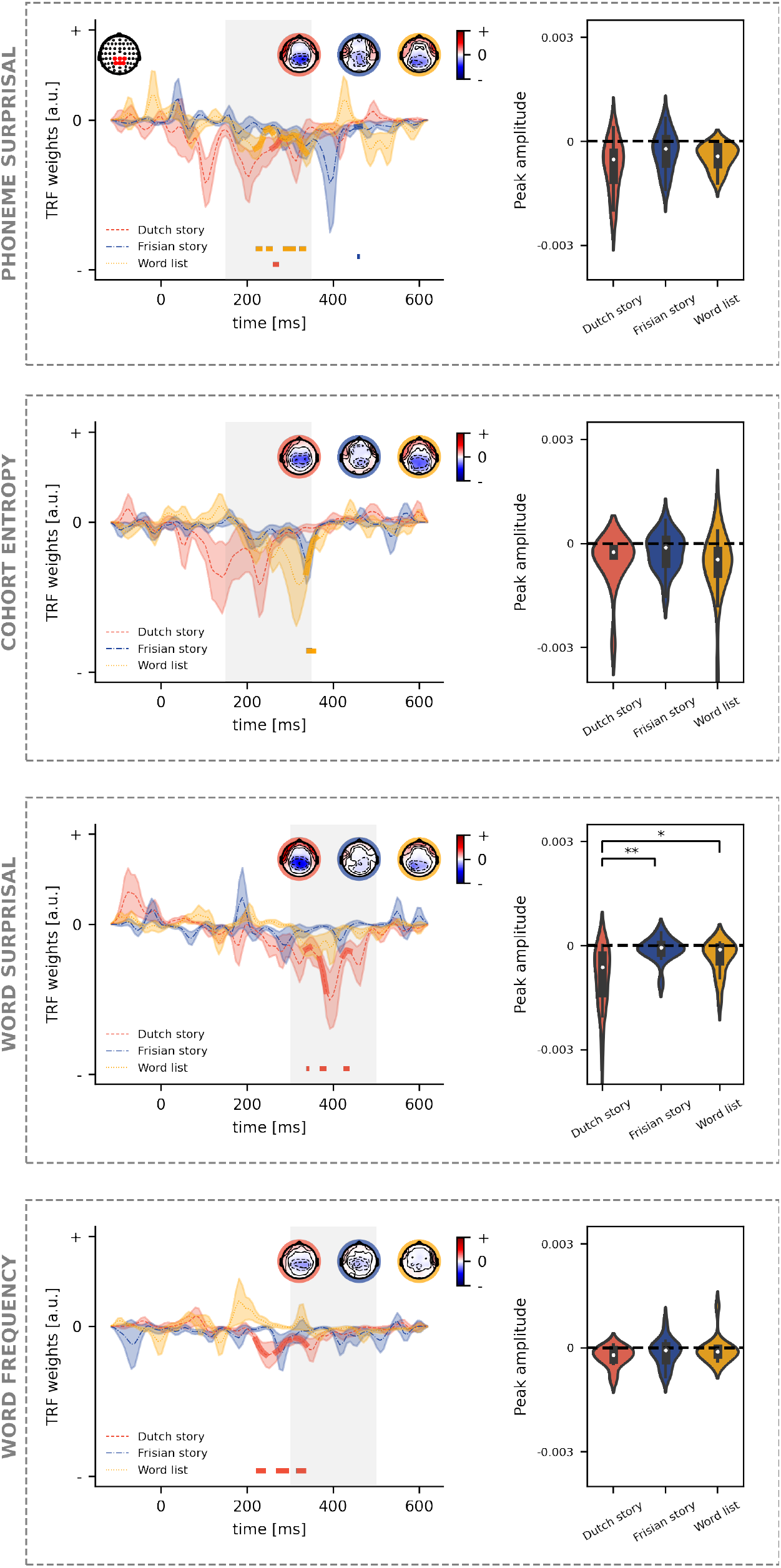
Neural response to language features. For each language feature, the TRFs in a central channel selection (shown as a left inset on the top panel) are depicted (left). The significance of the TRF is indicated by the thick lines in the corresponding colour and the horizontal lines below the TRF. The time region used for peak determination is marked by the vertical grey block. The associated peak topographies are visualized as insets on each panel for, respectively, the Dutch story (orange), Frisian story (blue) and word list (yellow). On the right side of each panel, the peak amplitudes are visualized. *(*: p < 0.05, **:p < 0.01, ***: p < 0.001)*

### 3.4 Neural response to language features

We identified whether the TRF was significantly different from zero (corrected for three comparisons).

For phoneme surprisal, we observe that all three speech materials show at least one cluster. For the Dutch story, we observed 1 significant cluster, from 259 to 275 ms. For the word list, 4 clusters were identified ranging from 220 to 236 ms, 243 to 259 ms, 282 to 314 ms and 322 to 337 ms. A small, marginal cluster at the boundary of the time region was identified for the Frisian story, i.e., from 455 to 462 ms. After peak determination, no differences in amplitude or latency between the three speech materials were found for phoneme surprisal evaluated in a central channel selection.

For cohort entropy, the *N* 250*_T RF_* of the TRF was less prominent than for phoneme surprisal. For the Dutch story as well as the Frisian story, no clusters were observed. One cluster was found for the word list ranging from 338 to 361 ms. Similar to phoneme surprisal, the differences between the peak amplitudes or latencies did not reach significance.

For word surprisal, the cluster-based permutation test identified three clusters ranging from 338 to 345 ms, 369 to 384 ms, and 423 to 439 ms. The TRF to word surprisal for the word list and the Frisian story did not show any clusters. Here, the amplitude difference reached significance: the *N* 400*_T RF_* amplitude for the Dutch story (median = -0.00063) is different from the Frisian story (median = -0.00005; u_(30)_ = 222, p *<* 0.001; corrected after Benjamini-Hochberg) as well as from the word list (median = -0.0001; u_(32)_ = 71, p = 0.024; corrected after Benjamini-Hochberg). Altogether these results suggest only a prominent *N* 400*_T RF_* is found in response to the Dutch story but not to the Frisian story or the word list. No differences were found regarding the latency of this *N* 400*_T RF_* peak.

For word frequency, we identified several clusters for the TRF for the Dutch story while no clusters were found for the Frisian story or the word list. For the Dutch story, we identified three clusters, ranging from 220 to 244 ms, 267 to 298 ms, and 314 to 338 ms. No differences were found regarding the amplitude or latency of the *N* 400*_T RF_* peak to word frequency.

## 4 Discussion

We observed that acoustic features were similarly tracked across the three different speech materials as seen by similar prediction accuracies of the acoustic model and similar patterns of neural responses (TRFs) for frontal channels. Regarding language tracking of speech, only the Dutch condition showed a significant improvement in prediction accuracy when language features are included on top of acoustic and (pre-)lexical features. For the TRFs in central channels, we observed different patterns for the different speech materials. For the Dutch fragments, all language features at word level and phoneme surprisal evoked a significant negative response. However, for word surprisal, only a prominent *N* 400*_T RF_* was observed in the TRF for the Dutch story, which significantly differed from the Frisian story and the word list.

Regarding the behavioral results, it is unsurprising that the Dutch story was better understood than the Frisian story and the word list. Although the study aimed to present an incomprehensible story with a similar syntax to the Dutch story, this was probably not achieved. The subjective ratings of speech understanding showed that some participants understood the Frisian story quite well. This is presumably because Frisian and Dutch are related languages which helps the listener to grasp the content of the story to some extent (Van Bezooijen and Gooskens, 2007). By getting an idea of the story’s content, the context could aid the word identification task, which could explain the higher word identification scores. A similar trend is seen in Broderick et al. (2020), the more the text is shuffled, the lower the performance on the word identification task. In sum, we could conclude that the content of the Dutch story is fully understood, while around 50% for the Frisian story and very limited for the content of the word list is captured by the participants. However, as a side note, the subjectively rated speech understanding is difficult to estimate to a story in a foreign language, i.e., it is not necessarily because someone thinks they understood 50% of the content that this is actually the case.

Previous studies suggest that the comprehension of speech is reflected by acoustic neural tracking. By manipulating the signal-to-noise ratio of the speech, Vanthornhout et al. (2018) showed that as speech comprehension increases, acoustic speech tracking increases as well. Additional evidence comes from the study by Etard and Reichenbach (2019). They investigated acoustic tracking to a comprehensible and foreign story and reported that acoustic tracking in the delta band encoded speech comprehension. However, we did not find a difference in acoustic tracking between the three speech materials. We attribute this to two substantial methodological differences: (1) speaker characteristics and (2) the choice of the frequency range. Firstly, Etard and Reichenbach (2019) did not keep the speaker constant while the speaker acoustics affect acoustic neural tracking. In the current study, we kept the speaker the same across the different speech materials to avoid this bias. Secondly, Etard and Reichenbach (2019) observed the effect in the delta band but not the theta band. Here, a broader frequency range is investigated. However, as EEG follows a 1/f-characteristics, a broader frequency range would follow similar patterns as the delta band. In line with our results, Kosem et al. (2022) and Verschueren et al. (2022) also did not see a link between acoustic tracking and speech comprehension. Altogether, this suggests that acoustic tracking is a necessity rather than a sufficient condition for speech comprehension.

In neural response patterns (the TRFs), no prominent differences were found between the three speech materials. This suggests that acoustic TRFs do not depend on whether or not the speech is understood. Often the P2 peak to acoustic features is associated with speech comprehension (Ding and Simon, 2012; Vanthornhout et al., 2019; Verschueren et al., 2022). However, here a P2 peak is found for all three speech materials. One hypothesis might be that the P2 peak is associated with speech segmentation but not necessarily comprehension. As the Frisian language follows a similar structure to Dutch, words could be segmented but not necessarily understood.

The results regarding prediction accuracy and TRF pattern of the (pre-)lexical features align with those observed for acoustic features. We did not observe a significant difference in prediction accuracy among the three speech materials. Likewise, the pattern in the TRF is fairly similar across the three speech material. Similar to the above, we speculate that this might indicate a successful speech segmentation in all three speech materials. Overall our results suggest that when someone listens attentively to speech in silence, neural tracking of acoustic and (pre-)lexical features is not sufficient to determine whether or not the listener understood the speech. A solution might be to investigate the neural tracking of language features.

We aimed to quantify language tracking, i.e., the extent the brain time-locks to the rhythm of language features, by investigating the increase in prediction accuracy when including language features and by investigating the presence of the *N* 250*_T RF_* and *N* 400*_T RF_* in the TRF. Only for the Dutch story, we observed that language features significantly improve the prediction accuracy compared to a model which only includes acoustic and (pre-)lexical features (Figure 4). This effect was prominent in a small cluster from central to left-lateralized channels. This finding is similar to the findings of Gillis et al. (2021) using the same language features. However, the effect was more prominent in the study of Gillis et al. (2021), which is likely due to the effect of more training data as impulsive features require more data (Mesik and Wojtczak, 2022). If the amount of data is small, the TRFs are noisier but have a similar pattern as those determined with more data (Mesik and Wojtczak, 2022). This justified investigating the TRFs for language features in more detail. Nevertheless, interpretation of the TRF without obtaining a significant improvement in prediction accuracy, as for the word list and Frisian story, should be done with care.

Regarding the TRFs at the phoneme level, we observed only one cluster for the Frisian story, which was situated around 450 ms for phoneme surprisal. The determination of phoneme surprisal is independent of previously uttered words and solely relies on the phoneme probability in the active cohort of words (unigram frequency of the word, independent of its context). As the phoneme frequencies are similar between Frisian and Dutch (Figure 1), we can expect that Dutch listeners try to form a word based on the uttered phonemes, which might explain the marginal negative response. The later negative peak, around 400 ms, to phoneme surprisal in the Frisian condition has a similar topography (not shown) as the *N* 250*_T RF_* response for the Dutch story. Presumably, the increase in latency indicates that more processing time is required to integrate the information of the new phoneme with the previously uttered phonemes to recognize a known word. For the Dutch story and the word list, the *N* 250*_T RF_* peak is observed, which is as expected. For these two speech materials, the active cohort can be estimated accurately, so a prediction of an upcoming phoneme can be made.

Cohort entropy also depends on the active cohort of words. We observed that the *N* 250*_T RF_* to cohort entropy is only visible for the word list but not for the Frisian story or the Dutch story. This result is surprising as we expected to observe a *N* 250*_T RF_* for the Dutch story as well. We speculate that this unexpected effect is due to the limited amount of data. (Gillis et al., 2021) showed that the added value of cohort entropy is smaller compared to the added value of phoneme surprisal (assessed on 45 minutes of speech). This lower effect size can explain why the results here for cohort entropy do not align with findings from previous studies.

Regarding the TRF at the word level, we observed that only the Dutch story obtains a significant *N* 400*_T RF_* for the word surprisal. The *N* 400*_T RF_* is thought to represent multiple processes, such as activation of the lexical item in memory and integration of the word in its context (for reviews, see Lau et al., 2008; Kutas and Federmeier, 2011). Therefore, it is not surprising that only an *N* 400*_T RF_* is observed for the Dutch story, as only for the Dutch story can a word be integrated into its context. As there is no meaningful context for the word list condition, a word cannot be integrated into its context, and therefore, it was expected and verified that there is no *N* 400*_T RF_* to word surprisal for the word list.

Although the Dutch listeners reported that they could understand around 50% of the Frisian story’s content, no *N* 400*_T RF_* was observed to word surprisal nor word frequency. The reasons are threefold. Firstly, the effect of an *N* 400*_T RF_* is not consistently present for all words as not all words are understood. Secondly, the 5-gram model captures the context of the four preceding words, which might be too limited as the participants only capture around half of the story’s content. Lastly, although the word frequencies are similar between Frisian and their translated items, it is not necessarily easier for the participant to map Frisian words to their Dutch translations. For example, *bern*, the Frisian word for children is unlikely to be mapped to its Dutch translation *kinderen* while for the Frisian word *twa* (two) is easier mapped to the Dutch translation *twee*.

Overall our results regarding the observed patterns in the TRFs to word surprisal converge with the findings of Di Liberto et al. (2021) and Broderick et al. (2020), who relied on semantic dissimilarity: the worse the listener can integrate a word in its context due to lower language proficiency or higher word scrambling, the less prominent the *N* 400*_T RF_* response. Di Liberto et al. (2021) tested participant groups with a different language proficiency starting from proficiency A, a level of language proficiency in which basic language knowledge is known. In the current study, we relied on Frisian, which was unfamiliar to the listeners. Unfortunately, we did not evaluate the proficiency of Frisian for the cohort of participants. Nevertheless, due to the closeness between Frisian and Dutch, some words could be understood. We estimate that the proficiency of the Flemish listeners listing to Frisian was below the language proficiency level A, explaining why no significant *N* 400*_T RF_* in our participant cohort while Di Liberto et al. (2021) did observe a significant *N* 400*_T RF_*, even for listeners with the lowest language proficiency.

For word frequency, negative clusters are found for the Dutch story but not for the word list or Frisian story. This does not align with our expectations: we expected to observe a *N* 400*_T RF_* for the word list as well, as word frequency is modeled independently of its context. Only a prominent *N* 400*_T RF_* was observed for word surprisal in the Dutch story, which significantly differed from the Frisian story and the word list. It is noticeable that the amplitude of *N* 400*_T RF_* negativity is smaller than word surprisal, which is in agreement with Gillis et al. (2021). Gillis et al. (2021) speculated that the response to word frequency might resemble the activation of a lexical item, while the response to word surprisal captures the activation of a lexical item and its integration into the context.

The current study’s caveats lie in determining language tracking. Firstly, the amount of data gathered for the Frisian story and word list condition is limited. This decreases the statistical power to obtain significant improvement in prediction accuracy and causes the TRFs to have a noisier pattern. However, we opted for a limited amount of data as listening to a word list, or incomprehensible story is less attractive, and attention might drift away from the speech stimulus. Secondly, we could further optimize language tracking by determining channel selection in a data-driven way. Here, the channel selection was determined based on Gillis et al. (2021). We believe that the channel selection should either be defined a priori, as was done in the current study, or should be determined in a data-driven, cross-validated way. However, as the data is limited, we opted for the first approach. Lastly, the current evaluation is performed at the population level. Due to the limited amount of data, the TRFs are noisy and, therefore, averaging across all participants improves the signal-to-noise ratio to allow interpretations of the TRF. We aimed to improve the signal-to-noise ratio using DSS across all participants using an unseen Dutch speech fragment. Therefore, it is possible that the application of DSS might bias the analysis. If more data is collected, one could relate the subjectively rated speech understanding to the increase in prediction accuracy or the pattern of the TRF of the language features. However, there is a trade-off between collecting more data and keeping participants motivated to attend an incomprehensible speech fragment for a longer time.

In sum, we only observed language tracking for the comprehensible story while not for the incomprehensible story or the word list. The most prominent effect was found for word surprisal: a prominent *N* 400*_T RF_* response was only present for the Dutch story, which significantly differed from the Frisian story or the word list. This corresponds with the hypothesis that the *N* 400*_T RF_* response to word surprisal resembles a word’s integration in its context. Overall, this indicates that language tracking is related to speech comprehension. Future research should focus on making this measure robust on a subject-specific level. If language tracking is suitable to evaluate speech comprehension at a subject-specific level, one could use it as an objective measure of speech comprehension, which would allow testing speech comprehension in populations that are currently difficult to test with behavioral tests, such as young children and persons with cognitive impairment.

## 5 Data Sharing Statement

The data (preprocessed data and code) that support the findings of this study can be made available upon request, so far as this is in agreement with privacy and ethical regulations.

## Acknowledgements

The authors would like to thank Marte De Jonghe for substantial help in collecting the data. Additionally, a big “thank you” goes to Babs Gezelle Meerburg en Ronald Noppers for their effort and help in creating the different speech materials. We acknowledge Hugo Van Hamme, Jelske Dijkstra, Henk van den Heuvel and Martijn Bentum for their aid in acquiring the language models used in this study. Lastly, we would like to thank all the participants for partaking in the study, without whom this study would not have been possible.

## 6 Supplementary Material

### 6.1 Comparison of feature distributions

We applied the Kolmogorov-Smirnov test to investigate whether the samples are drawn from the same underlying distribution (H0). Upon H0 rejection, one can assume that the distributions are drawn from different distributions. We here report the KS test statistic and the Holm-adjusted p-value (corrected for 3 comparisons; see Table S.1).

**Table S1:** KS test statistic and corrected p-value for each comparison of the distribution of values of a specific feature. (NL = Dutch story, FR = Frisian story, WL = word list)

| Phoneme surprisal |  |  |  | Cohort entropy |  |  |  |
| --- | --- | --- | --- | --- | --- | --- | --- |
|  | NL | FR | WL |  | NL | FR | WL |
| NL |  | 0.012<br>(p < 0.001) | 0.007<br>(p = 0.056) | NL |  | 0.022<br>(p < 0.001) | 0.007<br>(p = 0.038) |
| FR |  |  | 0.010<br>(p = 0.004) | FR |  |  | 0.015<br>(p < 0.001) |
| WL |  |  |  | WL |  |  |  |

| Word surprisal |  |  |  | Word frequency |  |  |  |
| --- | --- | --- | --- | --- | --- | --- | --- |
|  | NL | FR | WL |  | NL | FR | WL |
| NL |  | 0.002<br>(p = 1) | 0.004<br>(p = 1) | NL |  | 0.002<br>(p = 1) | 0.003<br>(p = 1) |
| FR |  |  | 0.004<br>(p = 1) | FR |  |  | 0.005<br>(p = 1) |
| WL |  |  |  | WL |  |  |  |

### 6.2 TRF across frequency bands: spectrogram and acoustic edges

**Figure S1:**
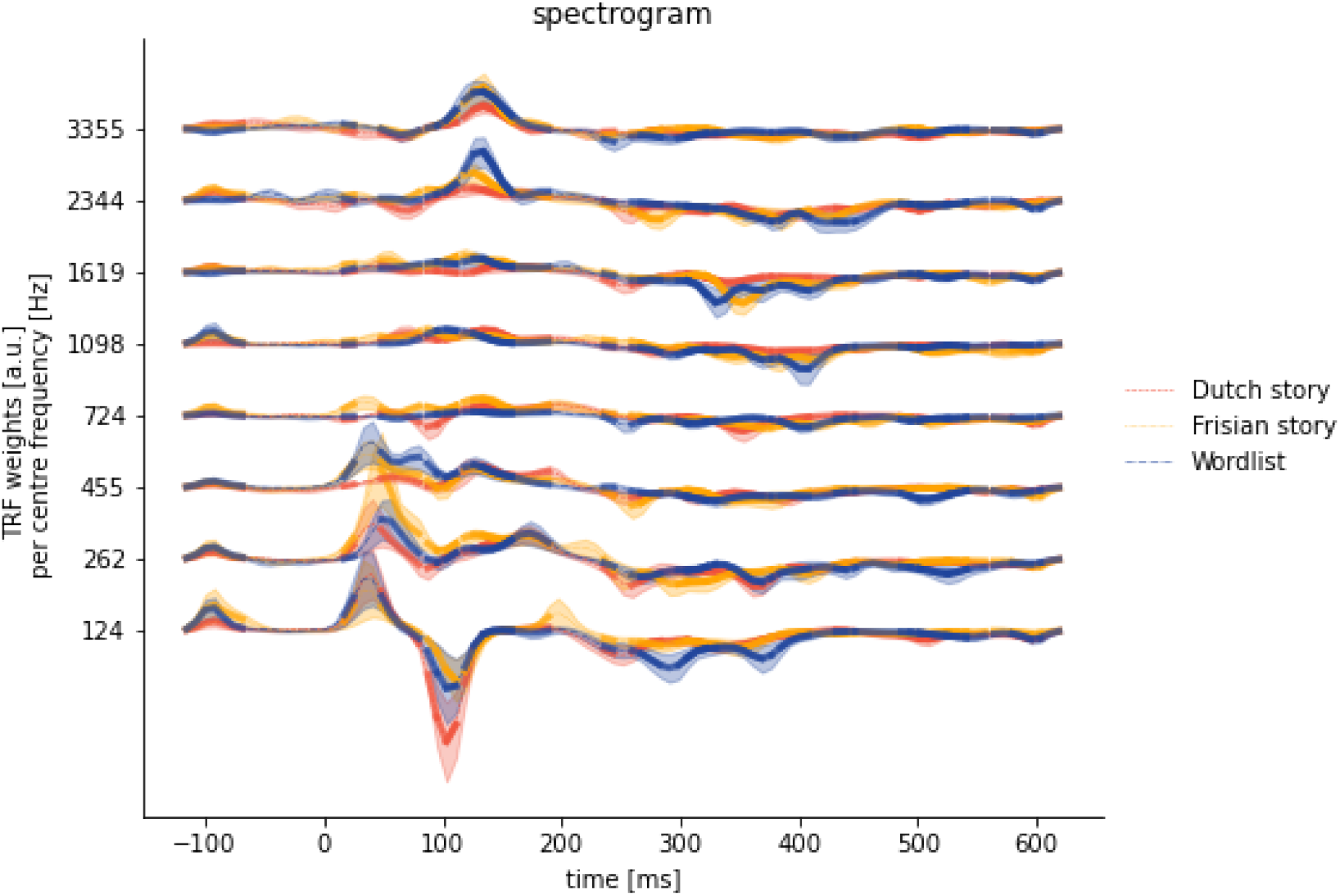
TRF for the spectrogram. The TRFs are visualized for each frequency band. The color indicates the speech material, respectively, the Dutch story (red), the Frisian story (yellow) and word list (blue).

**Figure S2:**
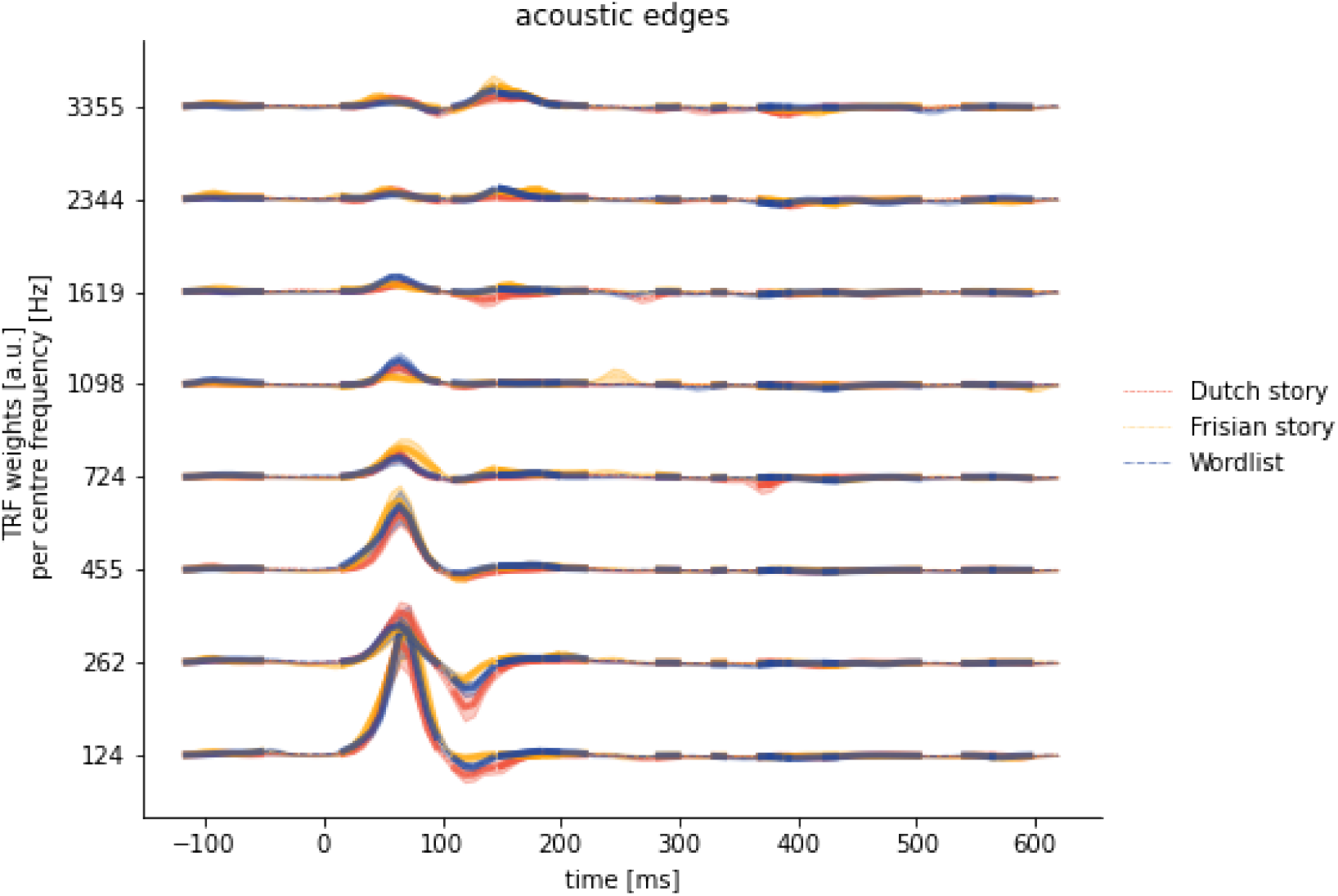
TRF for acoustic edges. The TRFs are visualized for each frequency band. The color indicates the speech material, respectively, the Dutch story (red), the Frisian story (yellow) and word list (blue).

